# Machine learning approaches to quantitively predict selectivity of compounds against hDAC1 and hDAC6 isoforms

**DOI:** 10.1101/2022.07.10.499476

**Authors:** Berna Dogan

## Abstract

The design of compounds selectively binding to specific isoforms of histone deacetylases (hDAC) is an ongoing research to prevent adverse side effects. Two of the most studied isoforms are hDAC1 and hDAC6 that are important targets to inhibit in various disease conditions. Here, various machine learning approaches were tested with the aim of developing models to predict the bioactivity and selectivity towards specific isoforms. Selectivity models were developed by directly training on the bioactivity differences of tested compounds against hDAC1 and hDAC6. Both classification and regression models were developed and compared to each other by using traditional evaluation metrics.

## Introduction

Histone deacetylases (hDAC) are enzymes catalysis the removal of acetyl group from proteins with acetylated lysine residues. The primary targets of hDACs are histone proteins surrounding the DNA and as such they are involved in regulating gene expression levels by altering DNA accessibility and chromatin structure.^1^ There are also variety of non-histone proteins targeted by hDACs such as transcription factors (E2F1, GATA1, NFkB, etc.), transcription suppressors (Yin Yang 1, YY1, Mad/Max), tumor suppressor gene p53, oncogene c-Myc.^2-4^ In addition, the functions of proteins such as cytoskeletal protein (α-tubulin),^5^ DNA helicase (WRN)^6^, heat shock protein (such as Hsp90)^7-8^ are controlled by hDAC enzymes. As such, hDAC enzymes are necessary for the normal body functions and disease conditions are observed when they are out of balance and/or overexpressed in the body.

In the human body, 18 different isoforms of hDAC enzymes have been identified until now. These enzymes are divided into four classes according to their structural similarities (homology), and their cellular localization.^9^ Class I enzymes include hDAC 1, 2, 3 and 8 isoforms and are usually found in the nucleus. Class II enzymes, on the other hand, are divided into IIa and IIb and though they are usually found in the cytoplasm, they can move to the nucleus when phosphorylated. Class IIa enzymes containing hDAC4, 5, 7, and 9 isoforms contain one catalytic domain, while class IIb enzymes consisting of hDAC6 and 10 have two catalytic domains. Class enzymes are also known as sirtuin proteins due to their homological similarity to silent information regulator 2 (silent information regulator 2) in yeast cells. Sirtuin proteins are also generally localized in the cytoplasm and they have seven isoforms (SIRT 1-7). Class IV enzymes containing the HDAC11 isoform are found both in the cytoplasm and in the nucleus, but it displays low homology with enzymes in other classes. Class I, II and IV enzymes function in a zinc (Zn^+2^) dependent manner while class III sirtuins function in Nicotinamide adenine dinucleotide (NAD^+^) dependent manner.

Molecules targeting hDAC enzymes have been in use for more than a decade especially in cancer treatment.^10-11^ However, approved hDAC inhibitors used today target more than one isoform of the enzyme, i.e. they are pan-inhibitors. Hence, these nonselective inhibitors could cause dose-dependent side effects in patients such as thrombocytopenia, cardiotoxicity, hematological toxicity, neutropenia, fatigue etc.^10, 12^ The discovery of molecules selective for hDAC isoforms, especially for hDAC6 enzyme, may allow for reduction of side effects as well as increase the effectivity of molecules specifically for diseases that depend on the function of hDAC6 enzyme. Though there is an intense research to discover hDAC6 specific inhibitors with many compounds being under clinical trials^12^ none of them has been approved yet.

Nowadays, molecular modeling techniques have become the basic components of many chemical, physical and biological studies. Computer-aided drug design methods are now routinely used to identify potential drug molecules and/or lead molecules, and they enable drug design stages to be carried out more rationally and quickly. There are varieties of techniques of simulation, virtual screening tools and algorithms for the identification of protein-ligand bindings profile including statistical approaches that employs binding data of existing molecules against specific targets for model developments. Qualitative structure-activity relationship (QSAR) studies was the first of these methods in which the biological activity of compounds were tried to be predicted by structural features of molecules. Though traditionally the underlying algorithm used in QSAR approaches has been based on multivariate linear and/or logistic regression methods^13-14^, other machine learning approaches such as decision tree based approaches, support vector machines, etc. started to be employed lately.^15-17^

Computational approaches have also utilized to predict the selectivity of compounds against target proteins.^13, 15-30^ In some studies, similarity searches were performed by using various descriptors^20^ and similarity coefficients such as Tanimoto.^21^ Many of these studies involve the development of individual QSAR or machine learning approaches were built for each target of interest and then selectivity profile was determined by comparison of predicted values. For instance, Tinivella et al. have developed accurate binary classification models for carbonic anhydrase isoforms II, IX and XII and determined selectivity profiles by the predicted active/inactive labels of compounds.^17^ With the increase in the available bioactivity data for various targets and isoforms, models trained directly on selectivity ratios or differences were started to be developed. Notably, Zhao et al. developed two-step QSAR models to first classify compounds as hDAC1 or hDAC6 selective and then predict their biological activity against these isoforms.^25^ Burggraaff et al. on the other hand recently built quantitative selectivity models to for adenosine receptors subtypes A_1_and A_2A_ adenosine receptors that were directly trained on the affinity differences of compounds for subtypes.^15^ With their selectivity window model, the authors not only accurately predicted the subtype selectivity of compounds but they also the degree of selectivity were estimated due to their usage of continuous models.

In this paper, various binary classification and continuous machine learning approaches were built to predict bioactivity as well as selectivity of hDAC1 and hDAC6 isoforms. These two isoforms were selected as they belong to difference classes and are the most studied isoforms of hDACs against which variety of compounds tested. Here, in addition to building bioactivity models for each isoform individually, selectivity models were also developed in which the differences between activity values were used for training as done by Burggraaff et al., with selectivity-window modeling. Several machine learning algorithms were evaluated and classification and regression models were also compared.

## Methods

### Dataset Preparation

The datasets for hDAC1 and hDAC6 were extracted from ChEMBL database (version 30). The target type was selected as single protein with target ChEMBL ID for hDAC1 as CHEMBL325 and for hDAC6 as CHEMBL1865. For both isoforms, the activity records were filtered to contain only binding assay with activity measured as IC_50_, EC_50_, K_i_ and K_D_ with certain data, i.e. standard relation set as “=“. The ChEMBL activity value, pChEMBL^31^ was used to standardize all the values from different types of activity measurement. As there were several activity measurements for some compounds, these duplicate records were preprocessed. Firstly, IC_50_ values are prioritized over other types, hence it was kept and activity records with other types were discarded. Then, the mean values were calculated for remaining activity records and this values were considered as the biological activity. At the end, hDAC1, 4493 unique compounds with biological activity data were collected. For hDAC6, 2972 unique compounds with biological activity data were gathered. For selectivity models, a dataset referred as hDAC1/6 dataset have been derived based on whether compounds have activity data for both isoforms. This dataset contains 1911 unique molecules with bioactivity data for both isoforms. In all datasets, activity values were considered in pChEMBL values.

The molecules in hDAC1 and hDAC6 datasets were categorized depending on whether they had bioactivity values for both isoforms and if they did, what were the activity differences. The compounds were categorized as *single-points* if they only have pChEMBL values for one of the isoforms. On the other hand, compounds with activity values for both isoforms were further categorized into five different classes depending on the activity and selectivity criteria as explained. The molecules were classified as *non-binders* if pChEMBL < 6.3 for both isoforms considered; as *hDAC6-selective* if pChEMBL ≥ 6.3 for hDAC6 with activity being 100-fold greater compared to hDAC1; as *hDAC1-selective* if pChEMBL ≥ 6.3 for hDAC1 with activity being 100-fold greater compared to hDAC6; as *dual-binders* if pChEMBL ≥ 6.3 for both isoforms with activity difference ≤ 10; as *semi-selective* if activity and selectivity differences did not fit any of the other categories. The labeled dataset were given in Additional Files 1-3.

### Generation and Testing of Molecular Descriptors

The compounds were described as 2D fingerprints using RDKit software. The following fingerprints were generated: MACCS^32^, Morgan^33^ (radius size of 3)and MAP4^34^ (radius size of 2) fingerprints. The feature based variation of Morgan fingerprints was also considered as a separate fingerprint and denoted as MorganF. For Morgan and MAP4 fingerprints different bit sizes; 1024 or 2048 were also tested. In total 7 different arrays of descriptors were generated for model testing.

### Machine Learning Algorithms

All the machine learning models developed using Python *scikit-learn* module. Initially, Lazy Predict library (https://github.com/shankarpandala/lazypredict) was used to evaluate all possible regression and classification models in the *scikit-learn* module. The datasets were divided into training and test set with 80:20 ratio. The statistical metrics such as balanced accuracy and F1 score for classification models; R^2^ and Root Mean Square Error (RMSE) for regression models were calculated for test set molecules. For models developed using the different descriptors used (2D fingerprints with different bit sizes), the statistical metrics were given in Supporting Tables S1-S8 for classification and regression models, respectively. Then, five of the models were further selected according to their performances based on the F1 score for classification and R^2^ for regression models as well as their methodology being different. For these five regression and classification models, 1024 bits Morgan fingerprints were used. The hyperparameters for chosen models were fine-tuned using Optuna software framework^35^ using recommended ranges for each algorithm in manuals, guides and previous studies^36-37^ and displayed in Table 2. To compare classification and regression models, the traditionally classification metrics were also applied to the output of regression models. For both classification and regression models, 10-fold cross validation sets were used for training. K-fold cross validation was used for regression, while for classification Stratified K-fold was used to approximately have same percentages of each target class, i.e. actives/inactives or selective/nonselective. To test the robustness of models, optimized models were subjected to 10-fold cross validation with different random splitting of cross validation sets.

## Results

### Datasets Exploration

The datasets were obtained from publicly available ChEMBL database (version 30). The collected bioactivity values given in different types, i.e. IC_50_, EC_50_, K_i_, K_d_, were standardized by considering pChEMBL values. Before building the models, the datasets of compounds were explored and labeled based on their bioactivity values as well as the differences between the values. Compounds were assigned as “Single-points” if there was bioactivity data for only one of the isoforms. If there were bioactivity data for the compound for both isoforms, then depending on pChEMBL values and the differences between them, it was categorized. The compound was labelled as “Non-binder” if pChEMBL < 6.3 for both isoforms while they were termed as “Dual-binder” if pChEMBL ≥ 6.3 for both isoforms and the ratio of the activity values ≤ 10.0 (i.e. pChEMBL(hDAC6) – pChEMBL(hDAC1) ≤ 1.0 and pChEMBL(hDAC6) – pChEMBL(hDAC1) ≥ -1.0). On the other hand, compounds were regarded as selective when the ratio of activity values ≥ 100.0. If the compound had pChEMBL ≥ 6.3 for hDAC1 and selectivity ratio ≥ 100.0, it was labelled as “hDAC1-selective”. If the compound had pChEMBL ≥ 6.3 for hDAC6 and selectivity ratio ≥ 100.0, it was labelled as “hDAC6-selective”. The remaining compounds with bioactivity values for both isoforms but cannot be categorized in any of the mentioned ones were termed as “Semi-selective”. Based on this categorization, 617 compounds in hDAC1/6 dataset have been identified as Semi-selective compounds. Table 1 display the characteristics of the labeled datasets. 59 compounds were identified as hDAC1-selective while 268 compounds were identified as hDAC6-selective. There were 612 compounds labeled as Dual-binders and 355 compounds labeled as Non-binders. In hDAC1 dataset, 2583 compounds were identified as Single-points since bioactivity measurements for only hDAC1 isoform was determined for these compounds. On the other hand, in hDAC6 dataset the number of compounds identified as Single-points were 1061. Figure 1 shows the distribution of compounds based on their activities in hDAC1 and hDAC6 datasets. It should be noted that hDAC1 dataset contained more than two times the number of compounds labeled as Single-points though the number of selective compounds for this isoform were considerably less such that hDAC6-selective compounds were more than four times larger than hDAC1-selective compounds. This could be due to the selectivity of molecules being more significant for hDAC6 isoform as explained this isoform is involved in many signaling pathway that could affect different diseases initiation and progression.

**Table 1.** Characteristics of datasets used for models development.

| Dataset | Number of Molecules | Depiction of Dataset | Activity (pChEMBL) |  |  |  |
| --- | --- | --- | --- | --- | --- | --- |
|  |  |  | isoform | Min | Median | Max |
| hDAC1 | 4493 | Molecules with activity values for hDAC1 | hDAC1 | 4.03 | 6.64 | 10.25 |
| hDAC6 | 2972 | Molecules with activity values for hDAC1 | hDAC6 | 4.00 | 7.00 | 10.10 |
| hDAC1/6 | 1294 | Molecules with activity values for both hDAC1 and hDAC6 (Dual-binder, Non-binder and Selective) | hDAC1* | 4.03 | 6.46 | 9.80 |
|  |  |  | hDAC6* | 4.03 | 7.20 | 10.00 |
| Semi-selective | 617 | Molecules with activity values for both hDAC1 and hDAC6 that do not fit into other categories (Semi-selective) | hDAC1* | 4.59 | 6.43 | 9.68 |
|  |  |  | hDAC6* | 4.79 | 7.12 | 9.15 |

\*Activity statistics calculated separately for hDAC1/6 and Semi-Selective dataset.

**Figure 1.**
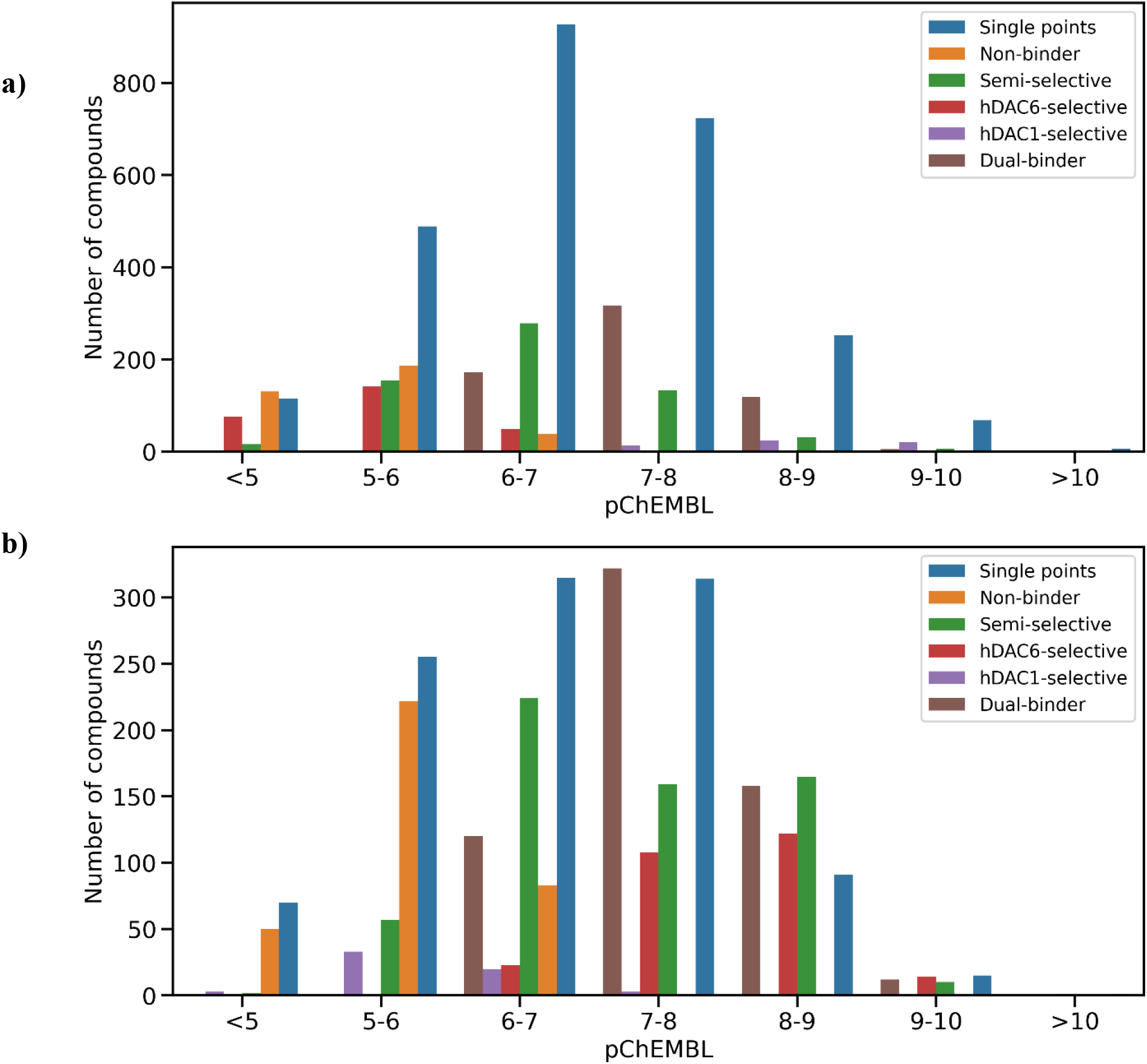
The bioactivity distribution and classification of compounds used in model development for isoforms a) hDAC1 b) hDAC6.

### Testing Machine Learning Algorithms

Bioactivity data for hDAC1 and hDAC6 isoforms were used to develop the classification and regression models. Models were developed to predict the bioactivity values of compounds for hDAC1 and hDAC6 isoform as well as to predict the selectivity of compounds for the specific isoform. Two different types of selectivity models were built: one with using hDAC1/6 dataset compounds, another with using both hDAC1/6 and Semi-selective dataset compounds.

Lazy predict library was utilized to evaluate the available machine learning algorithms in *scikit-learn* module of Python. For classification models, activity threshold was considered as pChEMBL ≥ 6.3. For both classification and regression models, the datasets were divided into training and test set by 80:20 ratio. 26 classification models were developed and evaluated based on their balanced accuracy (BA) and F1-score as displayed in Supporting Tables S1-S4. 24 regression models were tested and evaluated based on RMSE and R^2^ values as displayed in Supporting Tables S5-S8. Additionally, for each set all these models were built for 7 different fingerprint based descriptors that are 1024-bit and 2048-bit versions of Morgan, MorganF (feauture based Morgan) and MAP4 fingerprints as well as 166-bit sized MACCS fingerprints. The evaluation scores of models were not significantly affected by descriptor type and specifically the same machine learning algorithms were the ones with best evaluation scores for all descriptor types. Hence, Morgan fingerprints with 1024 bits were selected for further model development. Additionally, for both classification and regression, same five algorithms mentioned in the next section were always found to have best evaluation scores especially with Morgan 1024 bits descriptors. Hence, they were chosen to develop models whose hyperparameters was optimized for comparison.

The five machine learning algorithms chosen were namely: Random Forest (RF), Light Gradient Boosting Machine (LGBM), Extreme Gradient Boosting (XGB), K-Nearest Neighbors (KNN) and Support Vector Machines (SVM). Though histogram gradient boosting regressor algorithm were also among the models with highest evaluation scores, it was not preferred due to the longer time it requires to optimize this algorithm as it can not be parallelized. The hyperparameters for each of them were optimized using Optuna framework.^35^ The grid ranges for each hyperparameter of specific algorithm were applied as suggested in various studies^36-37^ as well as by searching various ranges and displayed in Table 2. In the optimization framework of Optuna, objective functions were generated such that evaluation metrics, F1-score for classification and R^2^ score for regression, were to be maximized. Models written in objective functions were trained by applying 10-fold cross validation iterators in which dataset considered was split into different sets and the evaluation metrics were averaged over these split sets. Then, study objects were created and objective functions were passed to the objects with optimization method. The number of trials for optimization were set as 1000 for LGBM, SVM, KNN and 100 for RF and SVM as the latest two algorithms were slower compared to others. After the trials were finished, the parameters which gave the best evaluation scores were detected and used to build optimized models.

**Table 2.**
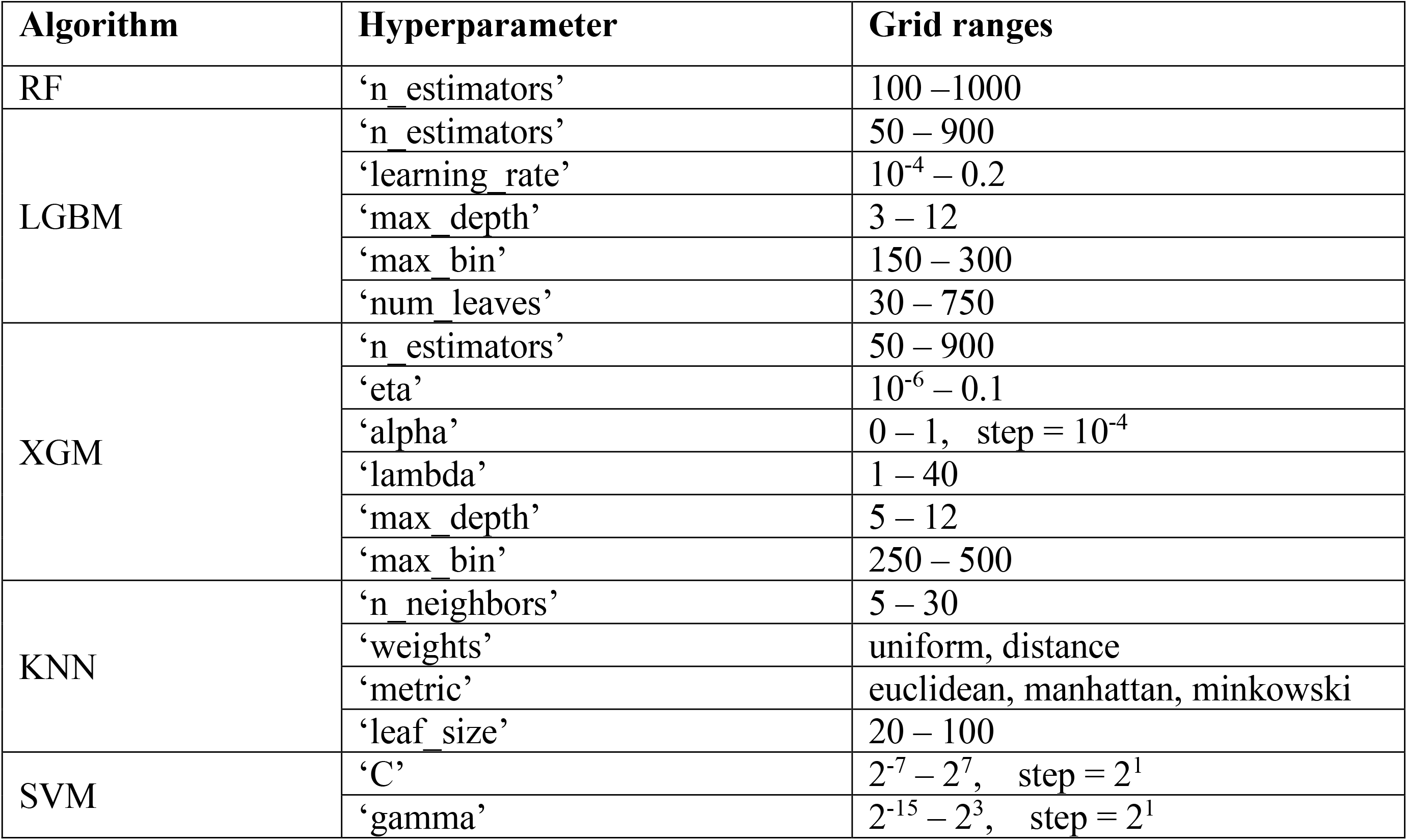
Selected machine learning algorithms and optimized hyperparameter with the grid ranges.

| Algorithm | Hyperparameter | Grid ranges |
| --- | --- | --- |
| RF | ‘n_estimators’ | 100 – 1000 |
| LGBM | ‘n_estimators’ | 50 – 900 |
| | ‘learning_rate’ | $10^{-4}$ – 0.2 |
|  | ‘max_depth’ | 3 – 12 |
|  | ‘max_bin’ | 150 – 300 |
|  | ‘num_leaves’ | 30 – 750 |
| XGM | ‘n_estimators’ | 50 – 900 |
| | ‘eta’ | $10^{-6}$ – 0.1 |
| | ‘alpha’ | 0 – 1, step = $10^{-4}$ |
|  | ‘lambda’ | 1 – 40 |
|  | ‘max_depth’ | 5 – 12 |
|  | ‘max_bin’ | 250 – 500 |
| KNN | ‘n_neighbors’ | 5 – 30 |
|  | ‘weights’ | uniform, distance |
|  | ‘metric’ | euclidean, manhattan, minkowski |
|  | ‘leaf_size’ | 20 – 100 |
| SVM | ‘C’ | $2^{-7}$ – $2^7$, step = $2^1$ |
| | ‘gamma’ | $2^{-15}$ – $2^3$, step = $2^1$ |

### Comparison of Classification and Regression Models

The performance of optimized models were evaluated by again 10-fold cross validation. Here, various other classification metrics such as accuracy, precision, sensitivity, specificity, F1-score, balanced accuracy (BA), Matthews Correlation Coefficient (MCC), Area Under the Receiver Operating Characteristic Curve (ROC AUC) were also calculated. The values were displayed in Table 3 for bioactivity prediction models with datasets hDAC1 and hDAC6. The evaluation parameters for selectivity models developed with hDAC1/6 dataset only and hDAC1/6 with Semi-selective compounds were given in Table 4. Here, for both classification and for the outcome of regression models, the selective window were determined such that compounds with selectivity difference larger than 2 were considered as selective. In other words, compounds considered selected where the ones whose pChEMBL(hDAC6) – pChEMBL(hDAC1) ≥ 2.0 or pChEMBL(hDAC6) – pChEMBL(hDAC1) ≤ -2.0. While the former ones with the differences higher than 2 were hDAC6 selective, the latter ones with differences less than -2 were hDAC1 selective compounds.

**Table 3.** Evaluation of optimized bioactivity models trained by 10-fold cross validation.

| **Classification bioactivity models for hDAC1** |  |  |  |  |  |
| --- | --- | --- | --- | --- | --- |
| *Metrics* | *RF* | *LGBM* | *XGB* | *KNN* | *SVM* |
| Accuracy | 0.84 ± 0.01 | 0.84 ± 0.02 | 0.85 ± 0.01 | 0.84 ± 0.02 | 0.85 ± 0.02 |
| Precision | 0.85 ± 0.01 | 0.86 ± 0.01 | 0.86 ± 0.01 | 0.87 ± 0.01 | 0.86 ± 0.02 |
| Sensitivity | 0.90 ± 0.02 | 0.89 ± 0.03 | 0.91 ± 0.02 | 0.87 ± 0.02 | 0.90 ± 0.02 |
| Specificity | 0.76 ± 0.03 | 0.77 ± 0.03 | 0.76 ± 0.02 | 0.79 ± 0.03 | 0.77 ± 0.03 |
| F1-score | 0.84 ± 0.01 | 0.84 ± 0.02 | 0.85 ± 0.01 | 0.84 ± 0.02 | 0.85 ± 0.02 |
| BA | 0.83 ± 0.02 | 0.83 ± 0.01 | 0.84 ± 0.01 | 0.83 ± 0.02 | 0.83 ± 0.02 |
| MCC | 0.66 ± 0.03 | 0.67 ± 0.03 | 0.68 ± 0.02 | 0.65 ± 0.03 | 0.68 ± 0.04 |
| ROC AUC | 0.83 ± 0.02 | 0.83 ± 0.01 | 0.84 ± 0.01 | 0.83 ± 0.02 | 0.83 ± 0.02 |
| **Regression bioactivity models for hDAC1** |  |  |  |  |  |
| R<sup>2</sup> | 0.69 ± 0.03 | 0.72 ± 0.03 | 0.72 ± 0.03 | 0.68 ± 0.03 | 0.73 ± 0.02 |
| Accuracy | 0.84 ± 0.02 | 0.84 ± 0.02 | 0.85 ± 0.02 | 0.84 ± 0.02 | 0.86 ± 0.02 |
| Precision | 0.84 ± 0.02 | 0.85 ± 0.02 | 0.86 ± 0.02 | 0.86 ± 0.02 | 0.87 ± 0.02 |
| Sensitivity | 0.91 ± 0.02 | 0.90 ± 0.02 | 0.90 ± 0.02 | 0.88 ± 0.02 | 0.91 ± 0.01 |
| Specificity | 0.73 ± 0.04 | 0.75 ± 0.03 | 0.77 ± 0.03 | 0.78 ± 0.03 | 0.78 ± 0.03 |
| F1-score | 0.84 ± 0.02 | 0.84 ± 0.02 | 0.85 ± 0.02 | 0.84 ± 0.02 | 0.86 ± 0.02 |
| BA | 0.82 ± 0.02 | 0.83 ± 0.02 | 0.83 ± 0.02 | 0.83 ± 0.02 | 0.84 ± 0.02 |
| MCC | 0.66 ± 0.04 | 0.66 ± 0.04 | 0.68 ± 0.04 | 0.66 ± 0.04 | 0.69 ± 0.03 |
| ROC AUC | 0.82 ± 0.02 | 0.83 ± 0.02 | 0.83 ± 0.02 | 0.83 ± 0.02 | 0.84 ± 0.02 |
| **Classification bioactivity models for hDAC6** |  |  |  |  |  |
| Accuracy | 0.86 ± 0.01 | 0.86 ± 0.01 | 0.86 ± 0.02 | 0.84 ± 0.02 | 0.87 ± 0.02 |
| Precision | 0.87 ± 0.01 | 0.88 ± 0.02 | 0.88 ± 0.02 | 0.87 ± 0.01 | 0.89 ± 0.02 |
| Sensitivity | 0.93 ± 0.01 | 0.93 ± 0.02 | 0.93 ± 0.01 | 0.92 ± 0.02 | 0.93 ± 0.02 |
| Specificity | 0.69 ± 0.04 | 0.70 ± 0.04 | 0.71 ± 0.05 | 0.68 ± 0.03 | 0.73 ± 0.04 |
| F1-score | 0.85 ± 0.01 | 0.86 ± 0.01 | 0.86 ± 0.02 | 0.84 ± 0.01 | 0.87 ± 0.02 |
| BA | 0.81 ± 0.02 | 0.82 ± 0.02 | 0.82 ± 0.03 | 0.80 ± 0.01 | 0.83 ± 0.02 |
| MCC | 0.65 ± 0.03 | 0.66 ± 0.03 | 0.67 ± 0.04 | 0.62 ± 0.03 | 0.68 ± 0.05 |
| ROC AUC | 0.81 ± 0.02 | 0.82 ± 0.02 | 0.82 ± 0.03 | 0.80 ± 0.01 | 0.83 ± 0.02 |
| **Regression bioactivity models for hDAC6** |  |  |  |  |  |
| R<sup>2</sup> | 0.69 ± 0.03 | 0.71 ± 0.03 | 0.70 ± 0.03 | 0.69 ± 0.03 | 0.65 ± 0.06 |
| Accuracy | 0.85 ± 0.02 | 0.86 ± 0.01 | 0.86 ± 0.02 | 0.85 ± 0.01 | 0.85 ± 0.02 |
| Precision | 0.87 ± 0.02 | 0.88 ± 0.02 | 0.88 ± 0.01 | 0.87 ± 0.01 | 0.86 ± 0.03 |
| Sensitivity | 0.93 ± 0.02 | 0.92 ± 0.01 | 0.93 ± 0.02 | 0.93 ± 0.02 | 0.92 ± 0.02 |
| Specificity | 0.68 ± 0.04 | 0.71 ± 0.04 | 0.70 ± 0.05 | 0.69 ± 0.04 | 0.67 ± 0.05 |
| F1-score | 0.85 ± 0.02 | 0.86 ± 0.01 | 0.86 ± 0.02 | 0.85 ± 0.01 | 0.84 ± 0.02 |
| BA | 0.80 ± 0.02 | 0.82 ± 0.02 | 0.82 ± 0.03 | 0.81 ± 0.02 | 0.80 ± 0.02 |
| MCC | 0.64 ± 0.04 | 0.66 ± 0.03 | 0.66 ± 0.05 | 0.64 ± 0.03 | 0.62 ± 0.04 |
| ROC AUC | 0.80 ± 0.02 | 0.82 ± 0.02 | 0.82 ± 0.03 | 0.81 ± 0.02 | 0.80 ± 0.02 |

**Table 4.** Evaluation of optimized selectivity models trained by 10-fold cross validation.

| **Classification selectivity models developed using hDAC1/6 dataset** |  |  |  |  |  |
| --- | --- | --- | --- | --- | --- |
| *Metrics* | *RF* | *LGBM* | *XGB* | *KNN* | *SVM* |
| Accuracy | 0.91 ± 0.03 | 0.92 ± 0.02 | 0.92 ± 0.03 | 0.92 ± 0.03 | 0.92 ± 0.02 |
| Precision | 0.88 ± 0.06 | 0.88 ± 0.04 | 0.88 ± 0.08 | 0.85 ± 0.07 | 0.85 ± 0.05 |
| Sensitivity | 0.76 ± 0.09 | 0.79 ± 0.09 | 0.78 ± 0.09 | 0.81 ± 0.08 | 0.80 ± 0.08 |
| Specificity | 0.96 ± 0.02 | 0.96 ± 0.02 | 0.96 ± 0.03 | 0.95 ± 0.03 | 0.96 ± 0.02 |
| F1-score | 0.91 ± 0.03 | 0.92 ± 0.02 | 0.91 ± 0.03 | 0.91 ± 0.03 | 0.92 ± 0.03 |
| BA | 0.86 ± 0.05 | 0.88 ± 0.04 | 0.87 ± 0.05 | 0.88 ± 0.04 | 0.88 ± 0.04 |
| MCC | 0.76 ± 0.08 | 0.78 ± 0.06 | 0.77 ± 0.09 | 0.77 ± 0.07 | 0.79 ± 0.07 |
| ROC AUC | 0.86 ± 0.05 | 0.88 ± 0.04 | 0.87 ± 0.05 | 0.88 ± 0.04 | 0.88 ± 0.04 |
| **Regression selectivity models developed using hDAC1/6 dataset** |  |  |  |  |  |
| R<sup>2</sup> | 0.72 ± 0.07 | 0.72 ± 0.05 | 0.73 ± 0.06 | 0.72 ± 0.08 | 0.73 ± 0.05 |
| Accuracy | 0.88 ± 0.03 | 0.88 ± 0.03 | 0.89 ± 0.03 | 0.90 ± 0.03 | 0.88 ± 0.04 |
| Precision | 0.93 ± 0.05 | 0.92 ± 0.06 | 0.91 ± 0.07 | 0.92 ± 0.08 | 0.95 ± 0.06 |
| Sensitivity | 0.58 ± 0.11 | 0.58 ± 0.11 | 0.62 ± 0.10 | 0.68 ± 0.07 | 0.54 ± 0.14 |
| Specificity | 0.99 ± 0.01 | 0.98 ± 0.01 | 0.98 ± 0.02 | 0.98 ± 0.02 | 0.99 ± 0.01 |
| F1-score | 0.87 ± 0.04 | 0.87 ± 0.04 | 0.88 ± 0.03 | 0.90 ± 0.03 | 0.86 ± 0.05 |
| BA | 0.78 ± 0.06 | 0.78 ± 0.06 | 0.80 ± 0.05 | 0.83 ± 0.04 | 0.77 ± 0.07 |
| MCC | 0.67 ± 0.10 | 0.67 ± 0.10 | 0.69 ± 0.09 | 0.74 ± 0.08 | 0.66 ± 0.11 |
| ROC AUC | 0.78 ± 0.06 | 0.78 ± 0.06 | 0.80 ± 0.05 | 0.83 ± 0.04 | 0.77 ± 0.07 |
| **Classification selectivity models developed using both hDAC1/6 and Semi-selective dataset** |  |  |  |  |  |
| Accuracy | 0.90 ± 0.03 | 0.91 ± 0.01 | 0.91 ± 0.02 | 0.90 ± 0.02 | 0.91 ± 0.02 |
| Precision | 0.77 ± 0.08 | 0.81 ± 0.07 | 0.82 ± 0.07 | 0.75 ± 0.07 | 0.76 ± 0.06 |
| Sensitivity | 0.62 ± 0.12 | 0.60 ± 0.09 | 0.61 ± 0.11 | 0.65 ± 0.10 | 0.68 ± 0.10 |
| Specificity | 0.96 ± 0.02 | 0.97 ± 0.01 | 0.97 ± 0.01 | 0.96 ± 0.01 | 0.96 ± 0.01 |
| F1-score | 0.90 ± 0.03 | 0.90 ± 0.02 | 0.91 ± 0.02 | 0.90 ± 0.02 | 0.91 ± 0.02 |
| BA | 0.79 ± 0.06 | 0.78 ± 0.04 | 0.79 ± 0.05 | 0.80 ± 0.05 | 0.82 ± 0.05 |
| MCC | 0.64 ± 0.10 | 0.64 ± 0.06 | 0.66 ± 0.09 | 0.64 ± 0.09 | 0.66 ± 0.08 |
| ROC AUC | 0.79 ± 0.06 | 0.78 ± 0.04 | 0.79 ± 0.05 | 0.80 ± 0.05 | 0.82 ± 0.05 |
| **Regression selectivity models developed using both hDAC1/6 and Semi-selective dataset** |  |  |  |  |  |
| R<sup>2</sup> | 0.72 ± 0.04 | 0.72 ± 0.04 | 0.74 ± 0.03 | 0.72 ± 0.05 | 0.74 ± 0.03 |
| Accuracy | 0.90 ± 0.02 | 0.89 ± 0.02 | 0.90 ± 0.02 | 0.90 ± 0.02 | 0.89 ± 0.02 |
| Precision | 0.83 ± 0.08 | 0.78 ± 0.07 | 0.78 ± 0.07 | 0.78 ± 0.08 | 0.81 ± 0.06 |
| Sensitivity | 0.53 ± 0.07 | 0.52 ± 0.09 | 0.55 ± 0.05 | 0.62 ± 0.10 | 0.50 ± 0.08 |
| Specificity | 0.98 ± 0.01 | 0.97 ± 0.01 | 0.97 ± 0.01 | 0.97 ± 0.01 | 0.98 ± 0.01 |
| F1-score | 0.89 ± 0.03 | 0.88 ± 0.03 | 0.89 ± 0.02 | 0.90 ± 0.03 | 0.88 ± 0.02 |
| BA | 0.75 ± 0.04 | 0.75 ± 0.05 | 0.76 ± 0.03 | 0.79 ± 0.05 | 0.74 ± 0.04 |
| MCC | 0.61 ± 0.08 | 0.58 ± 0.08 | 0.60 ± 0.06 | 0.64 ± 0.08 | 0.58 ± 0.07 |
| ROC AUC | 0.75 ± 0.04 | 0.75 ± 0.05 | 0.76 ± 0.03 | 0.79 ± 0.05 | 0.74 ± 0.04 |

## Supporting information

Additioal File 1 - hDAC1 compound dataset

Additioal File 2 - hDAC6 compound dataset

Additioal File 3 - hDAC 1 and 6 compound dataset

## Notes

### Competing Interest Statement

The authors have declared no competing interest.

## References

1. Gregoretti, I.; Lee, Y.-M.; Goodson, H. V., Molecular Evolution of the Histone Deacetylase Family: Functional Implications of Phylogenetic Analysis. J. Mol. Biol. 2004, 338 (1), 17–31.

2. Bolden, J. E.; Peart, M. J.; Johnstone, R. W., Anticancer activities of histone deacetylase inhibitors. Nat. Rev. Drug Discov. 2006, 5 (9), 769–84.

3. Elmallah, M. I. Y.; Micheau, O., Epigenetic Regulation of TRAIL Signaling: Implication for Cancer Therapy. Cancers 2019, 11 (6).

4. Glozak, M. A.; Sengupta, N.; Zhang, X.; Seto, E., Acetylation and deacetylation of non-histone proteins. Gene 2005, 363, 15–23.

5. Westermann, S.; Weber, K., Post-translational modifications regulate microtubule function. Nat. Rev. Mol. Cell Biol. 2003, 4 (12), 938–948.

6. Blander, G.; Zalle, N.; Daniely, Y.; Taplick, J.; Gray, M. D.; Oren, M., DNA damage-induced translocation of the Werner helicase is regulated by acetylation. J. Biol. Chem. 2002, 277 (52), 50934–50940.

7. Bali, P.; Pranpat, M.; Bradner, J.; Balasis, M.; Fiskus, W.; Guo, F.; Rocha, K.; Kumaraswamy, S.; Boyapalle, S.; Atadja, P., Inhibition of histone deacetylase 6 acetylates and disrupts the chaperone function of heat shock protein 90: a novel basis for antileukemia activity of histone deacetylase inhibitors. J. Biol. Chem. 2005, 280 (29), 26729–26734.

8. Kovacs, J. J.; Murphy, P. J. M.; Gaillard, S.; Zhao, X.; Wu, J.-T.; Nicchitta, C. V.; Yoshida, M.; Toft, D. O.; Pratt, W. B.; Yao, T.-P., HDAC6 regulates Hsp90 acetylation and chaperone-dependent activation of glucocorticoid receptor. Mol. Cell 2005, 18 (5), 601–607.

9. Kim, H.-J.; Bae, S.-C., Histone deacetylase inhibitors: molecular mechanisms of action and clinical trials as anti-cancer drugs. Am. J. Transl. Res. 2011, 3 (2), 166.

10. Li, T.; Zhang, C.; Hassan, S.; Liu, X.; Song, F.; Chen, K.; Zhang, W.; Yang, J., Histone deacetylase 6 in cancer. J. Hematol. Oncol. 2018, 11 (1), 111.

11. He, X.; Hui, Z.; Xu, L.; Bai, R.; Gao, Y.; Wang, Z.; Xie, T.; Ye, X.-Y., Medicinal chemistry updates of novel HDACs inhibitors (2020 to present). Eur. J. Med. Chem. 2022, 227, 113946.

12. Zhang, X. H.; Qin, M.; Wu, H. P.; Khamis, M. Y.; Li, Y. H.; Ma, L. Y.; Liu, H. M., A Review of Progress in Histone Deacetylase 6 Inhibitors Research: Structural Specificity and Functional Diversity. J. Med. Chem. 2021, 64 (3), 1362–1391.

13. Martin, E.; Mukherjee, P.; Sullivan, D.; Jansen, J., Profile-QSAR: a novel meta-QSAR method that combines activities across the kinase family to accurately predict affinity, selectivity, and cellular activity. J. Chem. Inf. Model. 2011, 51 (8), 1942–56.

14. Pham-The, H.; Casanola-Martin, G.; Dieguez-Santana, K.; Nguyen-Hai, N.; Ngoc, N. T.; Vu-Duc, L.; Le-Thi-Thu, H., Quantitative structure-activity relationship analysis and virtual screening studies for identifying HDAC2 inhibitors from known HDAC bioactive chemical libraries. SAR QSAR Environ. Res. 2017, 28 (3), 199–220.

15. Burggraaff, L.; van Vlijmen, H. W. T.; AP, I. J.; van Westen, G. J. P., Quantitative prediction of selectivity between the A1 and A2A adenosine receptors. J. Cheminform. 2020, 12 (1), 33.

16. Kurczab, R.; Canale, V.; Zajdel, P.; Bojarski, A. J., An Algorithm to Identify Target-Selective Ligands - A Case Study of 5-HT7/5-HT1A Receptor Selectivity. PLoS One 2016, 11 (6), e0156986.

17. Tinivella, A.; Pinzi, L.; Rastelli, G., Prediction of activity and selectivity profiles of human Carbonic Anhydrase inhibitors using machine learning classification models. J. Cheminform. 2021, 13 (1), 18.

18. Vogt, I.; Stumpfe, D.; Ahmed, H. E.; Bajorath, J., Methods for computer-aided chemical biology. Part 2: Evaluation of compound selectivity using 2D molecular fingerprints. Chem. Biol. Drug. Des. 2007, 70 (3), 195–205.

19. Stumpfe, D.; Geppert, H.; Bajorath, J., Methods for computer-aided chemical biology. Part 3: analysis of structure-selectivity relationships through single- or dual-step selectivity searching and Bayesian classification. Chem. Biol. Drug. Des. 2008, 71 (6), 518–28.

20. Ahmed, H. E.; Bajorath, J., Methods for computer-aided chemical biology. Part 5: rationalizing the selectivity of cathepsin inhibitors on the basis of molecular fragments and topological feature distributions. Chem. Biol. Drug Des. 2009, 74 (2), 129–41.

21. Peltason, L.; Hu, Y.; Bajorath, J., From structure-activity to structure-selectivity relationships: quantitative assessment, selectivity cliffs, and key compounds. ChemMedChem 2009, 4 (11), 1864–73.

22. Wassermann, A. M.; Geppert, H.; Bajorath, J., Searching for target-selective compounds using different combinations of multiclass support vector machine ranking methods, kernel functions, and fingerprint descriptors. J Chem Inf Model 2009, 49 (3), 582–92.

23. Ning, X.; Walters, M.; Karypis, G., Improved machine learning models for predicting selective compounds. J. Chem. Inf. Model. 2012, 52 (1), 38–50.

24. Ma, C.; Wang, L.; Yang, P.; Myint, K. Z.; Xie, X. Q., LiCABEDS II. Modeling of ligand selectivity for G-protein-coupled cannabinoid receptors. J. Chem. Inf. Model. 2013, 53 (1), 11–26.

25. Zhao, L.; Xiang, Y.; Song, J.; Zhang, Z., A novel two-step QSAR modeling work flow to predict selectivity and activity of HDAC inhibitors. Bioorg. Med. Chem. Lett. 2013, 23 (4), 929–33.

26. Li, Y.; Wang, L.; Liu, Z.; Li, C.; Xu, J.; Gu, Q.; Xu, J., Predicting selective liver X receptor beta agonists using multiple machine learning methods. Mol. Biosyst. 2015, 11 (5), 1241–50.

27. Montanari, F.; Zdrazil, B.; Digles, D.; Ecker, G. F., Selectivity profiling of BCRP versus P-gp inhibition: from automated collection of polypharmacology data to multi-label learning. J. Cheminform. 2016, 8, 7.

28. Hsu, K. C.; Liu, C. Y.; Lin, T. E.; Hsieh, J. H.; Sung, T. Y.; Tseng, H. J.; Yang, J. M.; Huang, W. J., Novel Class IIa-Selective Histone Deacetylase Inhibitors Discovered Using an in Silico Virtual Screening Approach. Sci. Rep. 2017, 7 (1), 3228.

29. Sciabola, S.; Stanton, R. V.; Wittkopp, S.; Wildman, S.; Moshinsky, D.; Potluri, S.; Xi, H., Predicting Kinase Selectivity Profiles Using Free-Wilson QSAR Analysis. J. Chem. Inf. Model. 2008, 48 (9), 1851–1867.

30. Zhang, J.; Han, B.; Wei, X.; Tan, C.; Chen, Y.; Jiang, Y., A two-step target binding and selectivity support vector machines approach for virtual screening of dopamine receptor subtype-selective ligands. PloS one 2012, 7 (6), e39076.

31. Papadatos, G.; Gaulton, A.; Hersey, A.; Overington, J. P., Activity, assay and target data curation and quality in the ChEMBL database. J. Comput. Aided Mol. Des. 2015, 29 (9), 885–896.

32. Durant, J. L.; Leland, B. A.; Henry, D. R.; Nourse, J. G., Reoptimization of MDL Keys for Use in Drug Discovery. J. Chem. Inf. Comput. Sci. 2002, 42 (6), 1273–1280.

33. Rogers, D.; Hahn, M., Extended-Connectivity Fingerprints. J. Chem. Inf. Model. 2010, 50 (5), 742–754.

34. Capecchi, A.; Probst, D.; Reymond, J. L., One molecular fingerprint to rule them all: drugs, biomolecules, and the metabolome. J. Cheminform. 2020, 12 (1), 43.

35. Akiba, T.; Sano, S.; Yanase, T.; Ohta, T.; Koyama, M., Optuna: A Next-generation Hyperparameter Optimization Framework. 1907.10902 2019.

36. Hsu, C.-W.; Chang, C.-C.; Lin, C.-J., A practical guide to support vector classification. Taipei, Taiwan: 2003.

37. Zhang, J.; Mucs, D.; Norinder, U.; Svensson, F., LightGBM: An Effective and Scalable Algorithm for Prediction of Chemical Toxicity-Application to the Tox21 and Mutagenicity Data Sets. J Chem Inf Model 2019, 59 (10), 4150–4158.

