## Supplementary Material for "Machine learning approaches to quantitively predict selectivity of compounds against hDAC1 and hDAC6 isoforms"

Berna Dogan

*Department of Biochemistry, School of Medicine, Bahcesehir University, Istanbul 34734, Turkey*

**Table S1. Comparison of classification algorithms for modeling bioactivity for hDAC1**

| Model | MACCS (166 bits) | | Morgan (1024 bits) | | MorganF (1024 bits) | | MAP4 (1024 bits) | | Morgan (2048 bits) | | MorganF (2048 bits) | | MAP4 (2048 bits) | |
|---|---|---|---|---|---|---|---|---|---|---|---|---|---|---|
| | BA | F1 | BA | F1 | BA | F1 | BA | F1 | BA | F1 | BA | F1 | BA | F1 |
| AdaBoostClassifier | 0.65 | 0.68 | 0.73 | 0.75 | 0.74 | 0.76 | 0.73 | 0.75 | 0.74 | 0.76 | 0.71 | 0.74 | 0.74 | 0.77 |
| BaggingClassifier | 0.80 | 0.81 | 0.80 | 0.81 | 0.80 | 0.81 | 0.79 | 0.80 | 0.80 | 0.81 | 0.81 | 0.82 | 0.79 | 0.80 |
| BernoulliNB | 0.60 | 0.62 | 0.74 | 0.74 | 0.74 | 0.74 | 0.62 | 0.63 | 0.75 | 0.76 | 0.74 | 0.73 | 0.64 | 0.65 |
| CalibratedClassifierCV | 0.68 | 0.70 | 0.72 | 0.74 | 0.72 | 0.75 | 0.67 | 0.70 | 0.73 | 0.76 | 0.74 | 0.77 | 0.71 | 0.75 |
| DecisionTreeClassifier | 0.77 | 0.78 | 0.74 | 0.76 | 0.77 | 0.79 | 0.71 | 0.72 | 0.75 | 0.77 | 0.77 | 0.79 | 0.69 | 0.71 |
| DummyClassifier | 0.51 | 0.53 | 0.51 | 0.53 | 0.51 | 0.53 | 0.52 | 0.54 | 0.51 | 0.55 | 0.51 | 0.55 | 0.51 | 0.55 |
| ExtraTreeClassifier | 0.75 | 0.76 | 0.76 | 0.77 | 0.76 | 0.78 | 0.65 | 0.67 | 0.75 | 0.76 | 0.75 | 0.77 | 0.67 | 0.70 |
| ExtraTreesClassifier | 0.80 | 0.81 | 0.84 | 0.86 | 0.82 | 0.83 | 0.80 | 0.82 | 0.83 | 0.84 | 0.83 | 0.85 | 0.81 | 0.83 |
| GaussianNB | 0.61 | 0.64 | 0.72 | 0.72 | 0.74 | 0.74 | 0.58 | 0.61 | 0.74 | 0.75 | 0.67 | 0.64 | 0.61 | 0.65 |
| KNeighborsClassifier | 0.78 | 0.80 | 0.79 | 0.80 | 0.79 | 0.80 | 0.76 | 0.78 | 0.78 | 0.79 | 0.78 | 0.80 | 0.78 | 0.81 |
| LabelPropagation | 0.75 | 0.75 | 0.53 | 0.32 | 0.55 | 0.39 | 0.52 | 0.29 | 0.53 | 0.27 | 0.56 | 0.36 | 0.51 | 0.23 |
| LabelSpreading | 0.75 | 0.75 | 0.53 | 0.32 | 0.55 | 0.39 | 0.52 | 0.29 | 0.53 | 0.27 | 0.56 | 0.36 | 0.51 | 0.23 |
| LGBMClassifier | 0.80 | 0.82 | 0.82 | 0.84 | 0.83 | 0.84 | 0.84 | 0.85 | 0.82 | 0.84 | 0.81 | 0.82 | 0.84 | 0.86 |
| LinearDiscriminantAnalysis | 0.68 | 0.70 | 0.76 | 0.77 | 0.78 | 0.79 | 0.72 | 0.74 | 0.75 | 0.76 | 0.70 | 0.72 | 0.68 | 0.70 |
| LinearSVC | 0.68 | 0.70 | 0.72 | 0.73 | 0.72 | 0.73 | 0.71 | 0.72 | 0.73 | 0.74 | 0.75 | 0.76 | 0.71 | 0.72 |
| NearestCentroid | 0.59 | 0.60 | 0.74 | 0.74 | 0.73 | 0.73 | 0.61 | 0.63 | 0.76 | 0.76 | 0.74 | 0.73 | 0.63 | 0.66 |
| NuSVC | 0.78 | 0.80 | 0.82 | 0.83 | 0.81 | 0.83 | 0.77 | 0.79 | 0.82 | 0.85 | 0.80 | 0.82 | 0.79 | 0.81 |
| PassiveAggressiveClassifier | 0.60 | 0.62 | 0.73 | 0.75 | 0.74 | 0.75 | 0.71 | 0.73 | 0.75 | 0.76 | 0.76 | 0.77 | 0.72 | 0.73 |
| Perceptron | 0.57 | 0.59 | 0.72 | 0.73 | 0.75 | 0.76 | 0.70 | 0.71 | 0.74 | 0.75 | 0.74 | 0.75 | 0.74 | 0.75 |
| QuadraticDiscriminantAnalysis | 0.61 | 0.63 | 0.64 | 0.67 | 0.68 | 0.71 | 0.59 | 0.61 | 0.71 | 0.65 | 0.65 | 0.70 | 0.51 | 0.51 |
| RandomForestClassifier | 0.79 | 0.81 | 0.83 | 0.85 | 0.82 | 0.83 | 0.81 | 0.82 | 0.84 | 0.86 | 0.82 | 0.84 | 0.81 | 0.84 |
| RidgeClassifier | 0.68 | 0.70 | 0.76 | 0.78 | 0.78 | 0.79 | 0.72 | 0.74 | 0.75 | 0.76 | 0.70 | 0.72 | 0.68 | 0.69 |
| RidgeClassifierCV | 0.68 | 0.70 | 0.76 | 0.78 | 0.78 | 0.79 | 0.72 | 0.75 | 0.76 | 0.77 | 0.74 | 0.75 | 0.68 | 0.70 |
| SGDClassifier | 0.65 | 0.66 | 0.72 | 0.74 | 0.74 | 0.75 | 0.70 | 0.71 | 0.73 | 0.76 | 0.72 | 0.75 | 0.72 | 0.74 |
| SVC | 0.76 | 0.78 | 0.81 | 0.83 | 0.81 | 0.83 | 0.76 | 0.79 | 0.82 | 0.84 | 0.80 | 0.82 | 0.77 | 0.80 |
| XGBClassifier | 0.80 | 0.81 | 0.82 | 0.83 | 0.82 | 0.83 | 0.83 | 0.84 | 0.82 | 0.84 | 0.82 | 0.84 | 0.82 | 0.84 |

BA represents Balanced Accuracy, F1 represents the F1 score.

**Table S2. Comparison of classification algorithms for modeling bioactivity for hDAC6**

| Model | MACCS (166 bits) | | Morgan (1024 bits) | | MorganF (1024 bits) | | MAP4 (1024 bits) | | Morgan (2048 bits) | | MorganF (2048 bits) | | MAP4 (2048 bits) | |
|---|---|---|---|---|---|---|---|---|---|---|---|---|---|---|
| | **BA** | **F1** | **BA** | **F1** | **BA** | **F1** | **BA** | **F1** | **BA** | **F1** | **BA** | **F1** | **BA** | **F1** |
| AdaBoostClassifier | 0.69 | 0.75 | 0.73 | 0.79 | 0.74 | 0.80 | 0.72 | 0.78 | 0.75 | 0.80 | 0.77 | 0.81 | 0.75 | 0.80 |
| BaggingClassifier | 0.78 | 0.82 | 0.79 | 0.83 | 0.80 | 0.84 | 0.77 | 0.82 | 0.84 | 0.86 | 0.80 | 0.83 | 0.77 | 0.81 |
| BernoulliNB | 0.68 | 0.73 | 0.75 | 0.78 | 0.74 | 0.78 | 0.68 | 0.72 | 0.78 | 0.79 | 0.77 | 0.79 | 0.63 | 0.66 |
| CalibratedClassifierCV | 0.71 | 0.78 | 0.69 | 0.76 | 0.70 | 0.77 | 0.61 | 0.69 | 0.71 | 0.78 | 0.72 | 0.79 | 0.64 | 0.73 |
| DecisionTreeClassifier | 0.76 | 0.79 | 0.76 | 0.81 | 0.77 | 0.81 | 0.72 | 0.76 | 0.78 | 0.81 | 0.77 | 0.80 | 0.75 | 0.78 |
| DummyClassifier | 0.49 | 0.56 | 0.49 | 0.56 | 0.49 | 0.56 | 0.53 | 0.59 | 0.49 | 0.56 | 0.49 | 0.56 | 0.49 | 0.56 |
| ExtraTreeClassifier | 0.76 | 0.79 | 0.77 | 0.80 | 0.75 | 0.79 | 0.65 | 0.70 | 0.75 | 0.78 | 0.80 | 0.82 | 0.63 | 0.69 |
| ExtraTreesClassifier | 0.79 | 0.82 | 0.80 | 0.85 | 0.81 | 0.85 | 0.73 | 0.80 | 0.83 | 0.86 | 0.80 | 0.84 | 0.75 | 0.81 |
| GaussianNB | 0.65 | 0.55 | 0.74 | 0.77 | 0.75 | 0.75 | 0.58 | 0.66 | 0.69 | 0.65 | 0.64 | 0.56 | 0.56 | 0.65 |
| KNeighborsClassifier | 0.77 | 0.82 | 0.77 | 0.81 | 0.79 | 0.82 | 0.74 | 0.80 | 0.76 | 0.81 | 0.74 | 0.78 | 0.74 | 0.80 |
| LabelPropagation | 0.77 | 0.77 | 0.52 | 0.24 | 0.56 | 0.35 | 0.52 | 0.25 | 0.52 | 0.22 | 0.55 | 0.32 | 0.52 | 0.21 |
| LabelSpreading | 0.77 | 0.77 | 0.52 | 0.24 | 0.56 | 0.35 | 0.52 | 0.25 | 0.52 | 0.22 | 0.55 | 0.32 | 0.52 | 0.21 |
| LGBMClassifier | 0.79 | 0.83 | 0.79 | 0.83 | 0.82 | 0.86 | 0.75 | 0.81 | 0.82 | 0.85 | 0.81 | 0.85 | 0.79 | 0.83 |
| LinearDiscriminantAnalysis | 0.71 | 0.78 | 0.75 | 0.80 | 0.78 | 0.81 | 0.70 | 0.74 | 0.61 | 0.63 | 0.68 | 0.70 | 0.61 | 0.64 |
| LinearSVC | 0.72 | 0.78 | 0.73 | 0.77 | 0.76 | 0.79 | 0.70 | 0.73 | 0.78 | 0.79 | 0.80 | 0.81 | 0.77 | 0.79 |
| NearestCentroid | 0.68 | 0.74 | 0.74 | 0.78 | 0.74 | 0.77 | 0.60 | 0.65 | 0.78 | 0.80 | 0.77 | 0.79 | 0.58 | 0.63 |
| NuSVC | 0.70 | 0.78 | 0.71 | 0.78 | 0.72 | 0.80 | 0.71 | 0.78 | 0.73 | 0.80 | 0.72 | 0.79 | 0.72 | 0.79 |
| PassiveAggressiveClassifier | 0.70 | 0.74 | 0.71 | 0.76 | 0.78 | 0.81 | 0.71 | 0.75 | 0.77 | 0.79 | 0.77 | 0.79 | 0.78 | 0.80 |
| Perceptron | 0.68 | 0.72 | 0.73 | 0.78 | 0.77 | 0.80 | 0.70 | 0.74 | 0.77 | 0.80 | 0.77 | 0.80 | 0.78 | 0.80 |
| QuadraticDiscriminantAnalysis | 0.58 | 0.47 | 0.53 | 0.59 | 0.59 | 0.67 | 0.51 | 0.57 | 0.61 | 0.44 | 0.68 | 0.60 | 0.54 | 0.29 |
| RandomForestClassifier | 0.79 | 0.83 | 0.80 | 0.85 | 0.81 | 0.86 | 0.75 | 0.81 | 0.82 | 0.85 | 0.80 | 0.84 | 0.78 | 0.83 |
| RidgeClassifier | 0.71 | 0.78 | 0.76 | 0.81 | 0.79 | 0.82 | 0.70 | 0.74 | 0.67 | 0.68 | 0.72 | 0.75 | 0.62 | 0.65 |
| RidgeClassifierCV | 0.71 | 0.78 | 0.77 | 0.82 | 0.79 | 0.82 | 0.71 | 0.75 | 0.75 | 0.77 | 0.76 | 0.78 | 0.69 | 0.71 |
| SGDClassifier | 0.71 | 0.76 | 0.72 | 0.78 | 0.74 | 0.79 | 0.67 | 0.73 | 0.72 | 0.79 | 0.73 | 0.80 | 0.74 | 0.79 |
| SVC | 0.75 | 0.81 | 0.77 | 0.83 | 0.78 | 0.83 | 0.69 | 0.76 | 0.75 | 0.82 | 0.76 | 0.82 | 0.71 | 0.79 |
| XGBClassifier | 0.79 | 0.83 | 0.79 | 0.84 | 0.83 | 0.87 | 0.75 | 0.80 | 0.82 | 0.85 | 0.81 | 0.85 | 0.79 | 0.83 |

BA represents Balanced Accuracy, F1 represents the F1 score.

**Table S3. Comparison of classification algorithms for modeling selectivity (Semi-selective compounds included)**

| Model | MACCS (166 bits) | | Morgan (1024 bits) | | MorganF (1024 bits) | | MAP4 (1024 bits) | | Morgan (2048 bits) | | MorganF (2048 bits) | | MAP4 (2048 bits) | |
|---|---|---|---|---|---|---|---|---|---|---|---|---|---|---|
| | **BA** | **F1** | **BA** | **F1** | **BA** | **F1** | **BA** | **F1** | **BA** | **F1** | **BA** | **F1** | **BA** | **F1** |
| AdaBoostClassifier | 0.65 | 0.83 | 0.71 | 0.85 | 0.67 | 0.84 | 0.75 | 0.88 | 0.69 | 0.85 | 0.70 | 0.85 | 0.73 | 0.88 |
| BaggingClassifier | 0.76 | 0.86 | 0.75 | 0.87 | 0.76 | 0.87 | 0.72 | 0.87 | 0.79 | 0.91 | 0.75 | 0.88 | 0.74 | 0.89 |
| BernoulliNB | 0.62 | 0.73 | 0.76 | 0.81 | 0.78 | 0.82 | 0.66 | 0.72 | 0.77 | 0.84 | 0.79 | 0.83 | 0.72 | 0.76 |
| CalibratedClassifierCV | 0.64 | 0.83 | 0.67 | 0.85 | 0.61 | 0.81 | 0.59 | 0.81 | 0.59 | 0.83 | 0.53 | 0.79 | 0.59 | 0.83 |
| DecisionTreeClassifier | 0.72 | 0.82 | 0.78 | 0.86 | 0.78 | 0.86 | 0.71 | 0.84 | 0.76 | 0.86 | 0.74 | 0.86 | 0.70 | 0.83 |
| DummyClassifier | 0.53 | 0.73 | 0.53 | 0.73 | 0.53 | 0.73 | 0.47 | 0.70 | 0.52 | 0.75 | 0.52 | 0.75 | 0.52 | 0.75 |
| ExtraTreeClassifier | 0.72 | 0.84 | 0.74 | 0.85 | 0.76 | 0.86 | 0.67 | 0.80 | 0.79 | 0.89 | 0.75 | 0.86 | 0.68 | 0.82 |
| ExtraTreesClassifier | 0.77 | 0.86 | 0.80 | 0.89 | 0.77 | 0.88 | 0.69 | 0.86 | 0.76 | 0.89 | 0.75 | 0.88 | 0.68 | 0.87 |
| GaussianNB | 0.54 | 0.19 | 0.69 | 0.69 | 0.66 | 0.68 | 0.58 | 0.74 | 0.68 | 0.78 | 0.68 | 0.82 | 0.72 | 0.84 |
| KNeighborsClassifier | 0.76 | 0.87 | 0.72 | 0.86 | 0.72 | 0.86 | 0.71 | 0.86 | 0.70 | 0.86 | 0.66 | 0.84 | 0.71 | 0.87 |
| LabelPropagation | 0.69 | 0.83 | 0.51 | 0.75 | 0.56 | 0.78 | 0.51 | 0.75 | 0.50 | 0.77 | 0.53 | 0.79 | 0.50 | 0.77 |
| LabelSpreading | 0.69 | 0.83 | 0.51 | 0.75 | 0.56 | 0.78 | 0.51 | 0.75 | 0.50 | 0.77 | 0.53 | 0.79 | 0.50 | 0.77 |
| LGBMClassifier | 0.77 | 0.86 | 0.82 | 0.89 | 0.78 | 0.87 | 0.76 | 0.89 | 0.77 | 0.89 | 0.75 | 0.87 | 0.76 | 0.90 |
| LinearDiscriminantAnalysis | 0.72 | 0.85 | 0.73 | 0.81 | 0.76 | 0.81 | 0.70 | 0.79 | 0.70 | 0.76 | 0.71 | 0.79 | 0.65 | 0.73 |
| LinearSVC | 0.74 | 0.86 | 0.73 | 0.81 | 0.74 | 0.82 | 0.69 | 0.81 | 0.73 | 0.82 | 0.77 | 0.83 | 0.77 | 0.85 |
| NearestCentroid | 0.65 | 0.69 | 0.78 | 0.83 | 0.79 | 0.83 | 0.59 | 0.70 | 0.76 | 0.85 | 0.76 | 0.84 | 0.67 | 0.74 |
| NuSVC | 0.66 | 0.78 | 0.73 | 0.82 | 0.77 | 0.84 | 0.71 | 0.82 | 0.75 | 0.84 | 0.78 | 0.85 | 0.78 | 0.88 |
| PassiveAggressiveClassifier | 0.73 | 0.82 | 0.75 | 0.83 | 0.74 | 0.83 | 0.72 | 0.83 | 0.76 | 0.84 | 0.76 | 0.85 | 0.74 | 0.86 |
| Perceptron | 0.63 | 0.62 | 0.53 | 0.76 | 0.57 | 0.79 | 0.50 | 0.74 | 0.58 | 0.38 | 0.56 | 0.50 | 0.55 | 0.20 |
| QuadraticDiscriminantAnalysis | 0.75 | 0.86 | 0.79 | 0.89 | 0.75 | 0.87 | 0.68 | 0.86 | 0.75 | 0.89 | 0.72 | 0.87 | 0.71 | 0.88 |
| RandomForestClassifier | 0.70 | 0.86 | 0.73 | 0.82 | 0.76 | 0.82 | 0.69 | 0.80 | 0.74 | 0.80 | 0.73 | 0.82 | 0.72 | 0.79 |
| RidgeClassifier | 0.70 | 0.86 | 0.74 | 0.83 | 0.75 | 0.83 | 0.70 | 0.82 | 0.77 | 0.82 | 0.75 | 0.85 | 0.71 | 0.82 |
| RidgeClassifierCV | 0.72 | 0.84 | 0.73 | 0.85 | 0.72 | 0.85 | 0.67 | 0.85 | 0.61 | 0.83 | 0.66 | 0.85 | 0.60 | 0.83 |
| SGDClassifier | 0.69 | 0.85 | 0.74 | 0.88 | 0.71 | 0.86 | 0.67 | 0.86 | 0.67 | 0.86 | 0.68 | 0.87 | 0.64 | 0.85 |
| SVC | 0.77 | 0.87 | 0.79 | 0.88 | 0.78 | 0.88 | 0.76 | 0.89 | 0.75 | 0.89 | 0.76 | 0.89 | 0.79 | 0.90 |
| XGBClassifier | 0.65 | 0.83 | 0.71 | 0.85 | 0.67 | 0.84 | 0.75 | 0.88 | 0.69 | 0.85 | 0.70 | 0.85 | 0.73 | 0.88 |

BA represents Balanced Accuracy, F1 represents the F1 score.

**Table S4. Comparison of classification algorithms for modeling selectivity (Semi-selective compounds not included)**

| **Model** | **MACCS (166 bits)** | | **Morgan (1024 bits)** | | **MorganF (1024 bits)** | | **MAP4 (1024 bits)** | | **Morgan (2048 bits)** | | **MorganF (2048 bits)** | | **MAP4 (2048 bits)** | |
|---|---|---|---|---|---|---|---|---|---|---|---|---|---|---|
| | **BA** | **F1** | **BA** | **F1** | **BA** | **F1** | **BA** | **F1** | **BA** | **F1** | **BA** | **F1** | **BA** | **F1** |
| AdaBoostClassifier | 0.78 | 0.85 | 0.82 | 0.87 | 0.82 | 0.86 | 0.82 | 0.87 | 0.77 | 0.86 | 0.80 | 0.87 | 0.79 | 0.87 |
| BaggingClassifier | 0.82 | 0.87 | 0.85 | 0.90 | 0.85 | 0.89 | 0.80 | 0.87 | 0.87 | 0.92 | 0.84 | 0.90 | 0.81 | 0.90 |
| BernoulliNB | 0.73 | 0.77 | 0.82 | 0.85 | 0.82 | 0.85 | 0.71 | 0.75 | 0.81 | 0.86 | 0.85 | 0.88 | 0.73 | 0.77 |
| CalibratedClassifierCV | 0.73 | 0.83 | 0.77 | 0.86 | 0.77 | 0.85 | 0.68 | 0.80 | 0.74 | 0.86 | 0.75 | 0.86 | 0.66 | 0.81 |
| DecisionTreeClassifier | 0.80 | 0.86 | 0.83 | 0.87 | 0.86 | 0.88 | 0.77 | 0.82 | 0.85 | 0.88 | 0.82 | 0.86 | 0.81 | 0.86 |
| DummyClassifier | 0.48 | 0.61 | 0.48 | 0.61 | 0.48 | 0.61 | 0.48 | 0.61 | 0.48 | 0.62 | 0.48 | 0.62 | 0.48 | 0.62 |
| ExtraTreeClassifier | 0.77 | 0.84 | 0.82 | 0.85 | 0.77 | 0.83 | 0.73 | 0.80 | 0.80 | 0.85 | 0.80 | 0.86 | 0.67 | 0.77 |
| ExtraTreesClassifier | 0.80 | 0.87 | 0.89 | 0.93 | 0.87 | 0.91 | 0.83 | 0.90 | 0.86 | 0.92 | 0.85 | 0.91 | 0.84 | 0.92 |
| GaussianNB | 0.54 | 0.25 | 0.77 | 0.75 | 0.69 | 0.7 | 0.66 | 0.76 | 0.63 | 0.75 | 0.67 | 0.78 | 0.60 | 0.73 |
| KNeighborsClassifier | 0.82 | 0.87 | 0.83 | 0.88 | 0.79 | 0.85 | 0.87 | 0.92 | 0.73 | 0.85 | 0.78 | 0.86 | 0.79 | 0.87 |
| LabelPropagation | 0.73 | 0.83 | 0.50 | 0.63 | 0.55 | 0.69 | 0.50 | 0.64 | 0.52 | 0.69 | 0.56 | 0.73 | 0.50 | 0.67 |
| LabelSpreading | 0.73 | 0.83 | 0.50 | 0.63 | 0.55 | 0.69 | 0.50 | 0.64 | 0.52 | 0.69 | 0.56 | 0.73 | 0.50 | 0.67 |
| LGBMClassifier | 0.80 | 0.87 | 0.89 | 0.92 | 0.87 | 0.91 | 0.87 | 0.92 | 0.87 | 0.92 | 0.84 | 0.90 | 0.82 | 0.90 |
| LinearDiscriminantAnalysis | 0.80 | 0.85 | 0.64 | 0.70 | 0.69 | 0.73 | 0.50 | 0.53 | 0.75 | 0.81 | 0.72 | 0.76 | 0.73 | 0.78 |
| LinearSVC | 0.80 | 0.86 | 0.85 | 0.87 | 0.81 | 0.84 | 0.80 | 0.84 | 0.88 | 0.90 | 0.85 | 0.86 | 0.83 | 0.86 |
| NearestCentroid | 0.72 | 0.73 | 0.82 | 0.86 | 0.82 | 0.85 | 0.64 | 0.73 | 0.81 | 0.87 | 0.86 | 0.88 | 0.64 | 0.73 |
| NuSVC | 0.67 | 0.79 | 0.61 | 0.75 | 0.64 | 0.78 | 0.58 | 0.73 | 0.66 | 0.81 | 0.65 | 0.80 | 0.62 | 0.78 |
| PassiveAggressiveClassifier | 0.73 | 0.81 | 0.83 | 0.87 | 0.82 | 0.84 | 0.80 | 0.84 | 0.89 | 0.92 | 0.84 | 0.87 | 0.85 | 0.88 |
| Perceptron | 0.76 | 0.83 | 0.84 | 0.88 | 0.83 | 0.86 | 0.81 | 0.86 | 0.81 | 0.87 | 0.84 | 0.86 | 0.86 | 0.91 |
| QuadraticDiscriminantAnalysis | 0.66 | 0.65 | 0.61 | 0.39 | 0.62 | 0.48 | 0.58 | 0.32 | 0.61 | 0.39 | 0.65 | 0.49 | 0.56 | 0.25 |
| RandomForestClassifier | 0.81 | 0.88 | 0.87 | 0.91 | 0.87 | 0.91 | 0.86 | 0.92 | 0.86 | 0.92 | 0.86 | 0.91 | 0.82 | 0.90 |
| RidgeClassifier | 0.79 | 0.85 | 0.69 | 0.75 | 0.76 | 0.81 | 0.66 | 0.72 | 0.76 | 0.82 | 0.79 | 0.83 | 0.75 | 0.81 |
| RidgeClassifierCV | 0.80 | 0.86 | 0.80 | 0.84 | 0.81 | 0.86 | 0.77 | 0.83 | 0.78 | 0.85 | 0.80 | 0.85 | 0.82 | 0.88 |
| SGDClassifier | 0.77 | 0.83 | 0.82 | 0.88 | 0.76 | 0.84 | 0.71 | 0.82 | 0.74 | 0.85 | 0.71 | 0.84 | 0.75 | 0.87 |
| SVC | 0.80 | 0.87 | 0.85 | 0.91 | 0.81 | 0.87 | 0.77 | 0.87 | 0.82 | 0.90 | 0.80 | 0.89 | 0.75 | 0.87 |
| XGBClassifier | 0.82 | 0.89 | 0.89 | 0.92 | 0.87 | 0.89 | 0.87 | 0.92 | 0.87 | 0.92 | 0.84 | 0.91 | 0.82 | 0.90 |

BA represents Balanced Accuracy, F1 represents the F1 score.

**Table S5. Comparison of regression algorithms for modeling bioactivity for hDAC1**

| Model | MACCS (166 bits) | | Morgan (1024 bits) | | MorganF (1024 bits) | | MAP4 (1024 bits) | | Morgan (2048 bits) | | MorganF (2048 bits) | | MAP4 (2048 bits) | |
|---|---|---|---|---|---|---|---|---|---|---|---|---|---|---|
| | $R^2$ | RMSE | $R^2$ | RMSE | $R^2$ | RMSE | $R^2$ | RMSE | $R^2$ | RMSE | $R^2$ | RMSE | $R^2$ | RMSE |
| AdaBoostRegressor | 0.22 | 1.02 | 0.22 | 1.03 | 0.19 | 1.04 | 0.38 | 0.90 | 0.14 | 1.04 | 0.17 | 1.02 | 0.44 | 0.84 |
| BaggingRegressor | 0.64 | 0.69 | 0.66 | 0.67 | 0.66 | 0.68 | 0.53 | 0.78 | 0.69 | 0.62 | 0.66 | 0.66 | 0.56 | 0.74 |
| BayesianRidge | 0.39 | 0.91 | 0.59 | 0.74 | 0.59 | 0.74 | 0.36 | 0.91 | 0.63 | 0.68 | 0.63 | 0.68 | 0.45 | 0.83 |
| DecisionTreeRegressor | 0.44 | 0.87 | 0.39 | 0.90 | 0.46 | 0.85 | 0.06 | 1.10 | 0.44 | 0.84 | 0.42 | 0.85 | 0.16 | 1.03 |
| ElasticNetCV | 0.39 | 0.90 | 0.58 | 0.75 | 0.61 | 0.73 | 0.34 | 0.92 | 0.61 | 0.70 | 0.64 | 0.68 | 0.41 | 0.86 |
| ExtraTreesRegressor | 0.45 | 0.86 | 0.46 | 0.85 | 0.54 | 0.79 | 0.58 | 0.74 | 0.50 | 0.79 | 0.47 | 0.82 | 0.63 | 0.68 |
| GammaRegressor | 0.27 | 0.99 | 0.55 | 0.78 | 0.56 | 0.77 | 0.36 | 0.91 | 0.60 | 0.71 | 0.60 | 0.71 | 0.00 | 1.12 |
| GeneralizedLinearRegressor | 0.28 | 0.99 | 0.55 | 0.77 | 0.56 | 0.77 | 0.36 | 0.91 | 0.60 | 0.71 | 0.60 | 0.70 | 0.45 | 0.83 |
| GradientBoostingRegressor | 0.46 | 0.85 | 0.52 | 0.80 | 0.55 | 0.78 | 0.58 | 0.74 | 0.51 | 0.78 | 0.50 | 0.79 | 0.61 | 0.70 |
| HistGradientBoostingRegressor | 0.64 | 0.69 | 0.68 | 0.65 | 0.70 | 0.64 | 0.66 | 0.67 | 0.70 | 0.62 | 0.67 | 0.64 | 0.68 | 0.63 |
| KNeighborsRegressor | 0.57 | 0.76 | 0.63 | 0.70 | 0.62 | 0.72 | 0.51 | 0.79 | 0.52 | 0.78 | 0.53 | 0.76 | 0.59 | 0.72 |
| LarsCV | 0.26 | 0.99 | 0.44 | 0.87 | 0.33 | 0.95 | 0.30 | 0.95 | 0.43 | 0.85 | 0.33 | 0.92 | 0.22 | 0.99 |
| LassoCV | 0.39 | 0.91 | 0.58 | 0.75 | 0.61 | 0.73 | 0.33 | 0.93 | 0.61 | 0.70 | 0.63 | 0.68 | 0.41 | 0.86 |
| LassoLarsCV | 0.39 | 0.90 | 0.57 | 0.76 | 0.60 | 0.73 | 0.31 | 0.94 | 0.57 | 0.74 | 0.59 | 0.72 | 0.38 | 0.88 |
| LassoLarsIC | 0.38 | 0.91 | 0.55 | 0.78 | 0.57 | 0.76 | 0.33 | 0.93 | 0.55 | 0.75 | 0.56 | 0.75 | 0.38 | 0.88 |
| LGBMRegressor | 0.64 | 0.69 | 0.68 | 0.65 | 0.70 | 0.64 | 0.67 | 0.65 | 0.70 | 0.62 | 0.67 | 0.64 | 0.68 | 0.63 |
| NuSVR | 0.62 | 0.72 | 0.67 | 0.66 | 0.68 | 0.66 | 0.52 | 0.79 | 0.63 | 0.68 | 0.63 | 0.69 | 0.54 | 0.76 |
| OrthogonalMatchingPursuit | 0.27 | 0.99 | 0.48 | 0.83 | 0.51 | 0.81 | 0.27 | 0.97 | 0.52 | 0.77 | 0.55 | 0.75 | 0.19 | 1.01 |
| OrthogonalMatchingPursuitCV | 0.27 | 0.99 | 0.48 | 0.84 | 0.51 | 0.81 | 0.21 | 1.01 | 0.51 | 0.78 | 0.55 | 0.75 | 0.22 | 0.99 |
| PoissonRegressor | 0.37 | 0.92 | 0.59 | 0.74 | 0.59 | 0.74 | 0.34 | 0.92 | 0.62 | 0.69 | 0.61 | 0.70 | 0.44 | 0.84 |
| RandomForestRegressor | 0.67 | 0.66 | 0.70 | 0.64 | 0.72 | 0.62 | 0.60 | 0.72 | 0.71 | 0.61 | 0.69 | 0.62 | 0.64 | 0.68 |
| SVR | 0.63 | 0.71 | 0.68 | 0.66 | 0.68 | 0.66 | 0.53 | 0.78 | 0.64 | 0.68 | 0.63 | 0.68 | 0.55 | 0.75 |
| TweedieRegressor | 0.28 | 0.99 | 0.55 | 0.77 | 0.56 | 0.77 | 0.36 | 0.91 | 0.60 | 0.71 | 0.60 | 0.70 | 0.45 | 0.83 |
| XGBRegressor | 0.67 | 0.67 | 0.67 | 0.66 | 0.68 | 0.65 | 0.61 | 0.71 | 0.66 | 0.65 | 0.67 | 0.65 | 0.63 | 0.68 |

**Table S6. Comparison of regression algorithms for modeling bioactivity for hDAC6**

| Model | MACCS (166 bits) | | Morgan (1024 bits) | | MorganF (1024 bits) | | MAP4 (1024 bits) | | Morgan (2048 bits) | | MorganF (2048 bits) | | MAP4 (2048 bits) | |
|---|---|---|---|---|---|---|---|---|---|---|---|---|---|---|
| | $R^2$ | RMSE | $R^2$ | RMSE | $R^2$ | RMSE | $R^2$ | RMSE | $R^2$ | RMSE | $R^2$ | RMSE | $R^2$ | RMSE |
| AdaBoostRegressor | 0.33 | 0.92 | 0.38 | 0.89 | 0.39 | 0.88 | 0.41 | 0.85 | 0.36 | 0.90 | 0.37 | 0.89 | 0.44 | 0.84 |
| BaggingRegressor | 0.65 | 0.67 | 0.68 | 0.63 | 0.69 | 0.62 | 0.58 | 0.72 | 0.62 | 0.69 | 0.65 | 0.67 | 0.59 | 0.72 |
| BayesianRidge | 0.45 | 0.84 | 0.65 | 0.67 | 0.64 | 0.67 | 0.39 | 0.86 | 0.59 | 0.72 | 0.61 | 0.70 | 0.46 | 0.83 |
| DecisionTreeRegressor | 0.43 | 0.85 | 0.40 | 0.87 | 0.43 | 0.85 | 0.20 | 0.99 | 0.47 | 0.82 | 0.41 | 0.86 | 0.22 | 0.99 |
| ElasticNetCV | 0.44 | 0.84 | 0.63 | 0.69 | 0.65 | 0.67 | 0.36 | 0.88 | 0.58 | 0.73 | 0.60 | 0.71 | 0.40 | 0.87 |
| ExtraTreesRegressor | 0.47 | 0.82 | 0.50 | 0.79 | 0.52 | 0.78 | 0.62 | 0.68 | 0.51 | 0.78 | 0.48 | 0.81 | 0.64 | 0.68 |
| GammaRegressor | 0.38 | 0.88 | 0.63 | 0.69 | 0.61 | 0.70 | 0.39 | 0.87 | 0.61 | 0.70 | 0.61 | 0.71 | 0.47 | 0.82 |
| GeneralizedLinearRegressor | 0.38 | 0.88 | 0.63 | 0.69 | 0.62 | 0.70 | 0.39 | 0.87 | 0.61 | 0.70 | 0.60 | 0.71 | 0.46 | 0.83 |
| GradientBoostingRegressor | 0.54 | 0.76 | 0.63 | 0.68 | 0.64 | 0.67 | 0.61 | 0.69 | 0.56 | 0.74 | 0.59 | 0.72 | 0.62 | 0.69 |
| HistGradientBoostingRegressor | 0.69 | 0.62 | 0.72 | 0.60 | 0.73 | 0.59 | 0.66 | 0.65 | 0.66 | 0.66 | 0.67 | 0.65 | 0.67 | 0.65 |
| KNeighborsRegressor | 0.59 | 0.72 | 0.63 | 0.69 | 0.59 | 0.72 | 0.49 | 0.79 | 0.54 | 0.76 | 0.53 | 0.77 | 0.51 | 0.79 |
| LarsCV | 0.40 | 0.87 | 0.57 | 0.74 | 0.49 | 0.80 | 0.32 | 0.91 | 0.51 | 0.78 | 0.44 | 0.84 | 0.27 | 0.96 |
| LassoCV | 0.44 | 0.84 | 0.62 | 0.69 | 0.65 | 0.67 | 0.36 | 0.89 | 0.58 | 0.73 | 0.60 | 0.71 | 0.40 | 0.87 |
| LassoLarsCV | 0.45 | 0.84 | 0.62 | 0.69 | 0.65 | 0.67 | 0.36 | 0.89 | 0.57 | 0.73 | 0.60 | 0.71 | 0.40 | 0.87 |
| LassoLarsIC | 0.44 | 0.84 | 0.56 | 0.75 | 0.57 | 0.73 | 0.35 | 0.89 | 0.52 | 0.78 | 0.57 | 0.73 | 0.35 | 0.91 |
| LGBMRegressor | 0.69 | 0.62 | 0.72 | 0.60 | 0.73 | 0.59 | 0.63 | 0.67 | 0.66 | 0.66 | 0.67 | 0.65 | 0.66 | 0.65 |
| NuSVR | 0.61 | 0.70 | 0.68 | 0.64 | 0.67 | 0.65 | 0.51 | 0.77 | 0.61 | 0.71 | 0.58 | 0.73 | 0.54 | 0.76 |
| OrthogonalMatchingPursuit | 0.36 | 0.90 | 0.57 | 0.74 | 0.56 | 0.74 | 0.21 | 0.98 | 0.40 | 0.87 | 0.47 | 0.81 | 0.19 | 1.01 |
| OrthogonalMatchingPursuitCV | 0.36 | 0.90 | 0.57 | 0.74 | 0.55 | 0.75 | 0.22 | 0.98 | 0.42 | 0.86 | 0.52 | 0.78 | 0.09 | 1.07 |
| PoissonRegressor | 0.44 | 0.84 | 0.63 | 0.68 | 0.63 | 0.68 | 0.34 | 0.90 | 0.53 | 0.77 | 0.58 | 0.73 | 0.41 | 0.87 |
| RandomForestRegressor | 0.68 | 0.64 | 0.72 | 0.60 | 0.73 | 0.59 | 0.63 | 0.68 | 0.65 | 0.67 | 0.68 | 0.64 | 0.64 | 0.68 |
| SVR | 0.62 | 0.69 | 0.68 | 0.63 | 0.67 | 0.64 | 0.51 | 0.77 | 0.61 | 0.70 | 0.58 | 0.73 | 0.55 | 0.76 |
| TweedieRegressor | 0.38 | 0.88 | 0.63 | 0.69 | 0.62 | 0.70 | 0.39 | 0.87 | 0.61 | 0.70 | 0.60 | 0.71 | 0.46 | 0.83 |
| XGBRegressor | 0.66 | 0.65 | 0.70 | 0.62 | 0.72 | 0.60 | 0.56 | 0.73 | 0.64 | 0.68 | 0.66 | 0.65 | 0.59 | 0.72 |

**Table S7. Comparison of regression algorithms for modeling selectivity (Semi-selective compounds included)**

| Model | MACCS (166 bits) | | Morgan (1024 bits) | | MorganF (1024 bits) | | MAP4 (1024 bits) | | Morgan (2048 bits) | | MorganF (2048 bits) | | MAP4 (2048 bits) | |
|---|---|---|---|---|---|---|---|---|---|---|---|---|---|---|
| | $R^2$ | RMSE | $R^2$ | RMSE | $R^2$ | RMSE | $R^2$ | RMSE | $R^2$ | RMSE | $R^2$ | RMSE | $R^2$ | RMSE |
| AdaBoostRegressor | 0.38 | 0.94 | 0.40 | 0.93 | 0.40 | 0.92 | 0.55 | 0.80 | 0.45 | 0.96 | 0.48 | 0.93 | 0.58 | 0.84 |
| BaggingRegressor | 0.60 | 0.76 | 0.64 | 0.71 | 0.65 | 0.71 | 0.58 | 0.77 | 0.75 | 0.65 | 0.74 | 0.65 | 0.69 | 0.71 |
| BayesianRidge | 0.42 | 0.91 | 0.62 | 0.74 | 0.63 | 0.73 | 0.41 | 0.92 | 0.73 | 0.67 | 0.68 | 0.73 | 0.56 | 0.85 |
| DecisionTreeRegressor | 0.48 | 0.86 | 0.47 | 0.87 | 0.43 | 0.90 | 0.32 | 0.98 | 0.64 | 0.77 | 0.67 | 0.75 | 0.40 | 1.00 |
| ElasticNetCV | 0.42 | 0.91 | 0.61 | 0.74 | 0.62 | 0.73 | 0.41 | 0.92 | 0.69 | 0.71 | 0.67 | 0.74 | 0.52 | 0.90 |
| ExtraTreesRegressor | 0.49 | 0.86 | 0.55 | 0.80 | 0.46 | 0.88 | 0.65 | 0.71 | 0.68 | 0.73 | 0.71 | 0.70 | 0.70 | 0.70 |
| GeneralizedLinearRegressor | 0.37 | 0.95 | 0.61 | 0.74 | 0.63 | 0.72 | -0.28 | 1.35 | 0.72 | 0.68 | 0.69 | 0.72 | 0.56 | 0.86 |
| GradientBoostingRegressor | 0.56 | 0.79 | 0.61 | 0.74 | 0.60 | 0.76 | 0.43 | 0.90 | 0.67 | 0.74 | 0.69 | 0.72 | 0.72 | 0.68 |
| HistGradientBoostingRegressor | 0.65 | 0.70 | 0.73 | 0.62 | 0.70 | 0.66 | 0.65 | 0.71 | 0.77 | 0.62 | 0.77 | 0.61 | 0.75 | 0.65 |
| KNeighborsRegressor | 0.57 | 0.78 | 0.63 | 0.72 | 0.61 | 0.75 | 0.61 | 0.74 | 0.65 | 0.77 | 0.64 | 0.77 | 0.61 | 0.81 |
| LarsCV | 0.37 | 0.94 | 0.32 | 0.98 | 0.47 | 0.87 | 0.28 | 1.01 | 0.56 | 0.85 | 0.47 | 0.93 | 0.35 | 1.04 |
| LassoCV | 0.42 | 0.91 | 0.61 | 0.74 | 0.62 | 0.74 | 0.40 | 0.92 | 0.69 | 0.72 | 0.67 | 0.74 | 0.50 | 0.91 |
| LassoLarsCV | 0.42 | 0.91 | 0.60 | 0.75 | 0.61 | 0.74 | 0.40 | 0.93 | 0.68 | 0.72 | 0.67 | 0.74 | 0.49 | 0.92 |
| LassoLarsIC | 0.42 | 0.91 | 0.47 | 0.87 | 0.54 | 0.81 | 0.33 | 0.97 | 0.00 | 1.29 | 0.00 | 1.29 | 0.00 | 1.29 |
| LGBMRegressor | 0.65 | 0.70 | 0.73 | 0.62 | 0.70 | 0.66 | 0.68 | 0.67 | 0.77 | 0.62 | 0.77 | 0.61 | 0.75 | 0.64 |
| NuSVR | 0.55 | 0.80 | 0.49 | 0.85 | 0.44 | 0.90 | 0.29 | 1.00 | 0.44 | 0.96 | 0.52 | 0.90 | 0.32 | 1.06 |
| OrthogonalMatchingPursuit | 0.60 | 0.76 | 0.66 | 0.69 | 0.67 | 0.69 | 0.54 | 0.81 | 0.67 | 0.74 | 0.67 | 0.74 | 0.53 | 0.89 |
| OrthogonalMatchingPursuitCV | 0.36 | 0.95 | 0.51 | 0.83 | 0.49 | 0.86 | 0.30 | 1.00 | 0.56 | 0.85 | 0.57 | 0.85 | 0.10 | 1.22 |
| RandomForestRegressor | 0.37 | 0.95 | 0.50 | 0.84 | 0.48 | 0.86 | 0.26 | 1.03 | 0.55 | 0.87 | 0.59 | 0.82 | 0.10 | 1.22 |
| SVR | 0.65 | 0.71 | 0.68 | 0.68 | 0.67 | 0.68 | 0.67 | 0.68 | 0.78 | 0.60 | 0.78 | 0.60 | 0.73 | 0.67 |
| TweedieRegressor | 0.60 | 0.76 | 0.66 | 0.69 | 0.67 | 0.69 | 0.54 | 0.81 | 0.67 | 0.74 | 0.68 | 0.73 | 0.54 | 0.88 |
| XGBRegressor | 0.37 | 0.95 | 0.61 | 0.74 | 0.63 | 0.72 | 0.43 | 0.90 | 0.72 | 0.68 | 0.69 | 0.72 | 0.56 | 0.86 |

| Table S8. Comparison of regression algorithms for modeling selectivity (Semi-selective compounds not included) | | | | | | | | | | | | | | |
|---|---|---|---|---|---|---|---|---|---|---|---|---|---|---|
| **Model** | **MACCS (166 bits)** | | **Morgan (1024 bits)** | | **MorganF (1024 bits)** | | **MAP4 (1024 bits)** | | **Morgan (2048 bits)** | | **MorganF (2048 bits)** | | **MAP4 (2048 bits)** | |
| | **$R^2$** | **RMSE** | **$R^2$** | **RMSE** | **$R^2$** | **RMSE** | **$R^2$** | **RMSE** | **$R^2$** | **RMSE** | **$R^2$** | **RMSE** | **$R^2$** | **RMSE** |
| AdaBoostRegressor | 0.46 | 0.95 | 0.50 | 0.92 | 0.41 | 0.99 | 0.52 | 0.89 | 0.37 | 0.97 | 0.31 | 1.02 | 0.58 | 0.80 |
| BaggingRegressor | 0.71 | 0.69 | 0.72 | 0.68 | 0.71 | 0.70 | 0.64 | 0.78 | 0.69 | 0.68 | 0.68 | 0.69 | 0.62 | 0.75 |
| BayesianRidge | 0.46 | 0.95 | 0.66 | 0.76 | 0.63 | 0.78 | 0.52 | 0.90 | 0.64 | 0.73 | 0.67 | 0.70 | 0.53 | 0.84 |
| DecisionTreeRegressor | 0.47 | 0.94 | 0.40 | 1.00 | 0.60 | 0.82 | 0.36 | 1.03 | 0.44 | 0.91 | 0.48 | 0.88 | 0.33 | 1.00 |
| ElasticNetCV | 0.46 | 0.95 | 0.61 | 0.81 | 0.65 | 0.77 | 0.42 | 0.98 | 0.65 | 0.72 | 0.67 | 0.70 | 0.41 | 0.94 |
| ExtraTreesRegressor | 0.52 | 0.90 | 0.52 | 0.90 | 0.63 | 0.78 | 0.68 | 0.73 | 0.46 | 0.89 | 0.49 | 0.87 | 0.68 | 0.69 |
| GeneralizedLinearRegressor | 0.40 | 1.00 | 0.66 | 0.75 | 0.64 | 0.77 | 0.50 | 0.92 | 0.68 | 0.69 | 0.69 | 0.68 | 0.54 | 0.82 |
| GradientBoostingRegressor | 0.62 | 0.80 | 0.71 | 0.70 | 0.63 | 0.79 | 0.70 | 0.70 | 0.66 | 0.71 | 0.66 | 0.72 | 0.71 | 0.66 |
| HistGradientBoostingRegressor | 0.73 | 0.67 | 0.75 | 0.65 | 0.70 | 0.71 | 0.73 | 0.67 | 0.72 | 0.65 | 0.72 | 0.64 | 0.71 | 0.66 |
| KNeighborsRegressor | 0.57 | 0.84 | 0.70 | 0.70 | 0.64 | 0.78 | 0.64 | 0.78 | 0.52 | 0.85 | 0.53 | 0.84 | 0.68 | 0.69 |
| LarsCV | 0.33 | 1.05 | 0.34 | 1.05 | 0.42 | 0.98 | 0.28 | 1.09 | 0.56 | 0.81 | 0.51 | 0.85 | 0.24 | 1.07 |
| LassoCV | 0.46 | 0.95 | 0.60 | 0.81 | 0.64 | 0.77 | 0.41 | 0.99 | 0.64 | 0.73 | 0.67 | 0.70 | 0.39 | 0.95 |
| LassoLarsCV | 0.45 | 0.95 | 0.60 | 0.82 | 0.64 | 0.78 | 0.41 | 0.99 | 0.65 | 0.73 | 0.67 | 0.71 | 0.41 | 0.94 |
| LassoLarsIC | 0.45 | 0.96 | 0.51 | 0.90 | 0.58 | 0.84 | 0.35 | 1.04 | -0.02 | 1.23 | -0.02 | 1.23 | -0.02 | 1.23 |
| LGBMRegressor | 0.73 | 0.67 | 0.75 | 0.65 | 0.70 | 0.71 | 0.72 | 0.69 | 0.72 | 0.65 | 0.72 | 0.64 | 0.70 | 0.67 |
| NuSVR | 0.68 | 0.74 | 0.55 | 0.87 | 0.52 | 0.90 | 0.40 | 1.00 | 0.22 | 1.08 | 0.36 | 0.98 | 0.47 | 0.89 |
| OrthogonalMatchingPursuit | 0.61 | 0.81 | 0.66 | 0.75 | 0.64 | 0.78 | 0.53 | 0.89 | 0.60 | 0.78 | 0.60 | 0.77 | 0.58 | 0.80 |
| OrthogonalMatchingPursuitCV | 0.44 | 0.96 | 0.40 | 1.00 | 0.51 | 0.90 | 0.27 | 1.11 | 0.39 | 0.96 | 0.45 | 0.90 | -0.24 | 1.36 |
| RandomForestRegressor | 0.44 | 0.96 | 0.49 | 0.92 | 0.51 | 0.90 | 0.28 | 1.10 | 0.51 | 0.86 | 0.51 | 0.85 | 0.19 | 1.10 |
| SVR | 0.73 | 0.67 | 0.75 | 0.65 | 0.72 | 0.68 | 0.68 | 0.74 | 0.71 | 0.66 | 0.71 | 0.66 | 0.67 | 0.70 |
| TweedieRegressor | 0.61 | 0.81 | 0.66 | 0.75 | 0.64 | 0.77 | 0.54 | 0.88 | 0.60 | 0.78 | 0.60 | 0.77 | 0.58 | 0.79 |
| XGBRegressor | 0.40 | 1.00 | 0.66 | 0.75 | 0.64 | 0.77 | 0.50 | 0.92 | 0.68 | 0.69 | 0.69 | 0.68 | 0.54 | 0.82 |
